# Robust estimates of cuticle conductance on stomatous leaf surfaces during the light induction of photosynthesis

**DOI:** 10.1101/2020.05.28.120030

**Authors:** Jun Tominaga, Joseph R. Stinziano, David T. Hanson

## Abstract

- Cuticle conductance (*g_cw_*) can bias calculations of intercellular CO_2_ concentration inside the leaf (*C_i_*) when stomatal conductance (*g_sw_*) is small.
- We examined how the light induction of photosynthesis impacts calculations by directly measuring *C_i_* along with standard gas exchange in sunflower and tobacco leaves.
- When photosynthesis was induced from dark to saturating light (1200 μmol m^−2^ s^−1^ PAR) the calculated *C_i_* was significantly larger than measured *C_i_* and the difference decreased as *g_sw_* increased. This difference could lead to over-estimation of rubisco deactivation by limited CO_2_ supply during early induction of photosynthesis. However, only small differences in *C_i_* were observed during the induction from shade (50 μmol m^−2^ s^−1^ PAR) because *g_sw_* was sufficiently large. The induction from dark also allowed robust estimations of *g_cw_* when combined with direct *C_i_* measurements. These *g_cw_* estimates succeeded in correcting the calculation, suggesting that the cuticle was the major source of error.
- Despite a technical restriction to amphi-stomatous leaves, the presented technique has a potential to provide insights into the cuticle conductance on intact stomatous leaf surfaces.

## Introduction

Since Moss and Rawlins (1963) first calculated intercellular CO_2_ concentration inside leaves (*C_i_*) from the gas exchange of single leaves, the technique has become standard practice in gas exchange studies. The calculations became gained utility when Farquhar et al. (1980) developed a photosynthesis model that uses *C_i_* to estimate substrate availability for ribulose-1,5-bisphosphate (RuBP) carboxylase oxygenase (rubisco). Based on a simple principle that rubisco carboxylation is limited by either of the two substrates, RuBP or CO_2_, the model predicts gas exchange of leaves as well as extrapolates underlying photosynthetic biochemistry (von Caemmerer, 2013). Accordingly, the calculations are the foundation of modern understanding of leaf carbon and water exchange.

However, calculations of *C_i_* have been questioned because uncertainties exist in assumptions (Hanson et al., 2016). *C_i_* is calculated from the relationship between water vapor exiting and CO_2_ entering the leaf by assuming both gasses only diffuse through stomata (Moss & Rawlins, 1963). In fact, water also diffuses through the cuticle. By including the cuticular transpiration (or cuticle conductance: *g_cw_*) in stomatal transpiration (or stomatal conductance: *g_sw_*), the calculation can overestimate the CO_2_ that has entered through stomata. Correction for cuticle effects is difficult as standard gas exchange measurements cannot distinguish cuticular and stomatal transpirations. Therefore, *g_cw_* is generally measured in isolated cuticles (Kerstiens, 1996; Schuster et al., 2017). The calculations also assume lateral uniformity in physicochemical properties over the leaf surface. Any heterogeneity, such as non-uniform stomatal apertures (Terashima, 1992; West et al., 2005), potentially bias the calculations as well (Terashima et al., 1988; Buckley, 1997; Meyer & Genty, 1998).

The accuracy of *C_i_* can be validated by directly measuring *C_i_* along with normal gas exchange measurements (Sharkey et al., 1982; Boyer & Kawamitsu, 2011; Tominaga & Kawamitsu, 2015a). In previous studies, amphi-stomatous leaves were put between an open gas exchange system and a closed system. The CO_2_ concentration inside the closed system is measured after it equilibrates with the *C_i_* while gas exchange is measured on the other side. Indeed, Sharkey et al. (1982) found that the measured *C_i_* was in close agreement to the value calculated on the other side when stomatal conductance was sufficiently large. Using a similar system, however, the over-estimation of calculated *C_i_* has been demonstrated by a large discrepancy from the measured *C_i_* when *g_sw_* is small (Boyer, 2015a; Tominaga & Kawamitsu, 2015b; Tominaga et al., 2018).

*C_i_* is critical for understanding how photosynthesis approaches a steady state during light induction. Within the first minute after a shift from low to high irradiance photosynthesis can be limited by the slow regeneration of RuBP while the Calvin Benson Cycle enzymes are activated (Sassenrath-Cole & Pearcy, 1992). Much of the rest of induction may be (co)-limited by stomatal opening and activation of rubisco. Upon illumination *C_i_* is depleted because stomatal opening is generally slower than activation of photosynthetic enzymes, thereby limiting rubisco carboxylation (Kirshbaum & Pearcy, 1988; Pearcy et al., 1996; Lawson and Blatt, 2014; McAusland et al., 2016; Dean et al., 2018). Meanwhile, rubisco is activated by sequential binding of CO_2_ and Mg^2+^ to the active site (Portis, 2003). This process is inhibited by a range of sugar-phosphates bound to the active site in darkness, and promoted by rubisco activase which releases these ligands in light. The reduction of *C_i_* also hinders this activation process, thereby limiting rubisco carboxylation indirectly (Mott & Woodrow, 1993). Therefore, initial *g_sw_* at the onset of light induction as well as the rapidity of stomatal opening have a large impact on gas exchange kinetics (Wachendorf & Küppers, 2017; Vialet-Chabrand et al., 2017). Both rapid stomatal regulation (Wang et al., 2014; Lawson & Blatt, 2014; McAusland et al., 2016) and rubisco activation (Yamori et al., 2012; Carmo-Silva & Salvucci, 2013; Soleh et al., 2016; Taylor and Long, 2017) can play significant roles and are novel targets for improving photosynthesis.

Because the initial *g_sw_* tends to be small in low irradiance or darkness the *C_i_* may be miscalculated during photosynthetic induction. Here we examine how light induction experiments impact the standard calculations by directly measuring *C_i_* along with gas exchange in leaves of sunflower (*Heliunthus annuus* L.) and tobacco (*Nicotiana tabacum* L.). Our results show when cuticle conductance matters and how it affects data interpretation. Together, we propose a novel technique to estimate intact cuticle conductance using the dual gas exchange system.

## Materials and methods

### Plant material

Tobacco (*N. tabacum* L. cv. Samsun) and sunflower (*H. annuus* L. cv. Mammoth Russian) plants, 9–11 weeks old and 5–7 weeks old respectively, were used. The tobacco plants were grown in 2.5-L plastic pots, and the sunflower plants were grown in 3.8-L plastic pots during December 2018 to April 2019 under ambient CO_2_ at 24.2/19.9 °C day/night temperature; each pot contained a soil mixture (Metro-Mix^®^ 360; Sun Gro Horticulture, Agawam, MA, USA) in a greenhouse located at the University of New Mexico [35.08°N, 106.62°W, 1587 m.a.s.l]. The plants were automatically watered three times each day and the pots were manually saturated with nutrient solutions containing 357 ppm N (Peters Professional^®^ 20-20-20 General Purpose; ICL Specialty Fertilizers, Summerville, SC, USA) and 195 ppm chelated Fe (Southern Agricultural Insecticides, Inc., Rubonia, FL, USA) twice and once weekly, respectively. Daylength in the greenhouse was extended to 15 h with LED lamps (VRPRx PLUS; Fluence Bioengineering, Inc., Austin, TX, USA) to prevent early flowering. All experiments with these plants were conducted with fully expanded upper leaves.

### Direct measurement system incorporated into LI-6800

The direct *C_i_* measurement system has previously been incorporated into a commercial open gas exchange system LI-6400 (LI-COR, Lincoln, NE, USA) and reported in detail elsewhere (Tominaga & Kawamitsu, 2015a). We applied a similar system for a new LI-6800. Instead of customizing a chamber, the manufactured LI-6800 chambers could be slightly modified to include the direct system (Fig. S1). In the present experiments, a large 6 x 6 cm chamber (6800-13; LI-COR) was used to maximize measurement precision, but the following system is applicable to the other chambers (e.g., Fluorometer chamber; 6800-01A; LI-COR). The chamber bottom-half was detached from the manifold and rotated 180°. Then, the flexible bellows originally attached to the bottom chamber were replaced with a solid adapter. The manifold to the bottom chamber was closed by a blanking plate with a rubber plate in between so that air only flows through the top chamber (i.e., open system). All experimental leaves were large enough to cover the entire chamber area, so the top and bottom chambers were divided by the leaf. Two miniature impeller pumps were connected in parallel to the adapter (inlet and outlet) of the rotated bottom chamber, and smoothly circulated the air to an infrared gas analyzer (IRGA; LI-7000; LI-COR) in a loop. When a sample leaf closed the loop, a water jacketed glass tube (condenser) maintained the dew point slightly lower than the room temperature (checked by the LI-7000), preventing condensation in the IRGA. In the closed loop, CO_2_ equilibrated with that inside the stomatal pores (*C_i(m)_*) was continuously measured while gas exchange through the opposite side of same leaf area was measured with the open system, which automatically calculated *C_i_* (*C_i(c)_*). Accordingly, both *C_i(m)_* and *C_i(c)_* were obtained simultaneously for the same leaf area. We configured LI-7000 to the differential mode by leading the air to the reference cell from the LI-6800 reference port (located at the backside of the sensor head) at a constant flow rate of 400 μmol s^−1^. The values of CO_2_ and water vapor concentrations in the reference air measured by the LI-6800 was exported to the LI-7000 whereas the sample CO_2_ concentration measured by the LI-7000 (i.e. *C_i(m)_*) was exported to and recorded by the LI-6800 along with other gas exchange parameters. By sharing the reference gas and data between the two systems, measurements were accurate and in sync. Plumbing of the closed loop was made mostly with copper tubing to avoid diffusion leaks. The advanced polymer gaskets (6568-512; LI-COR) and a large leaf area to edge ratio minimized the diffusion leaks such that any leaks were too small to be detected for the empty chamber in our experimental conditions, and no corrections were performed.

### Light induction

Plants were taken in the morning from the greenhouse to the laboratory. There, whole plants were acclimated in the dark for at least 1 h. Leaves were then clamped and illuminated by the chamber equipped with a large light source (6800-03; LI-COR). We set three levels of irradiances with white LEDs: 50, 200, and 1200 μmol m^−2^ s^−1^ photosynthetically active radiation (PAR) with the spectrum shown in Fig. S2. Measurements were automatically logged every 10 s until reaching a steady state. In all irradiances, full induction usually took less than 1.5 h and 2 h for tobacco and sunflower leaves, respectively. In addition to the induction from the dark, leaves were also subjected to saturating light (1200 μmol m^−2^ s^−1^ PAR) after being acclimated in shade (50 μmol m^−2^ s^−1^ PAR). Throughout the experiments, incoming air with a flow rate of 700 μmol s^−1^, air temperature of 25 °C, water vapor of 10 mmol mol^−1^ and CO_2_ concentration of 400 μmol mol^−1^ were held constant to avoid transient artefacts. Leaf temperature increased instantaneously upon illumination and was decreased throughout the induction by cooling from transpiration as stomata opened. Leaf temperature ranges varied among inductions: in tobacco, 24.5 ± 0.2□25.1 ± 0.1 °C (50 PAR), 24.4 ± 0.1□25.3 ± 0.1 °C (200 PAR), and 26.0 ± 0.1□27.4 ± 0.2°C (1200 PAR), and in sunflower, 24.0 ± 0.1□24.7 ± 0.1 °C (50 PAR), 24.1 ± 0.2□25.3 ± 0.2°C (200 PAR), and 25.8 ± 0.6 26.9 ± 0.4°C (1200 PAR), respectively.

### Calculations

The intercellular CO_2_ concentration inside the leaf (*C_i_*) was calculated from the gas exchange measurements on the adaxial leaf side, according to von Caemmerer and Farquhar (1981) as:

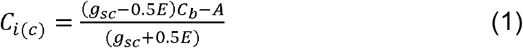

where *C_b_* is the CO_2_ concentration out of the leaf in the boundary layer (mol mol^−1^), *A* and *E* are the CO_2_ flux (mol CO_2_ m^−2^ s^−1^) and water vapor flux (mol H_2_O m^−2^ s^−1^), respectively, *g_sc_* is the stomatal conductance of CO_2_ (mol CO_2_ m^−2^ s^−1^). Given that *C_b_, A*, and *E* are measured variables, *g_sc_* is the critical parameter that determines validity of *C_i(c)_* (though measurement precision of *C_b_, A*, and *E* affect the calculation). Then, *g_sc_* would be expressed as:

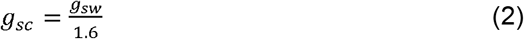

where *g_sw_* is the stomatal conductance of water vapor (mol H_2_O m^−2^ s^−1^) and 1.6 is the ratio of diffusivities of CO_2_ and water vapor in air. Then, *g_sw_* was calculated as:

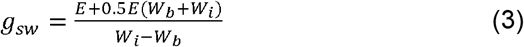

where *W_i_* and *W_b_* are the water vapor concentrations inside the leaf (mol mol^−1^) and out of the leaf in the boundary layer (mol mol^−1^), respectively. The *W_i_* was assumed to be saturated at the leaf temperature. However, the *g_sw_* includes cuticle conductance of water vapor (*g_cw_*) as:

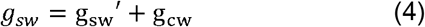

where *g_sw_’* is the strict sense of stomatal conductance of water vapor, which is only based on water vapor passing through the stomatal pore, and is in parallel with cuticle conductance (Jarvis, 1971).

Instead of estimating from *g_sw_* (Eq. (3)), we also calculated *g_sc_* from the directly measured *C_i_* (*C_i(m)_*) (Boyer and Kawamitsu, 2011) as:

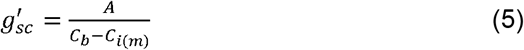

where *g_sc_’* was directly calculated from the measured CO_2_ gradients across the adaxial leaf surface.

According to the manufacturer’s calibration, boundary layer conductance of CO_2_ (1.5 CO_2_ mol m^−2^ s^−1^) and that of water vapor (2.1 mol H_2_O m^−2^ s^−1^) were estimated from the constant fan speed of 14,000 rpm, and used for calculating the *C_b_* and *W_b_*, respectively. A summary of parameters referred to within the text is shown with accompanying units in Table 1.

**Table 1.**
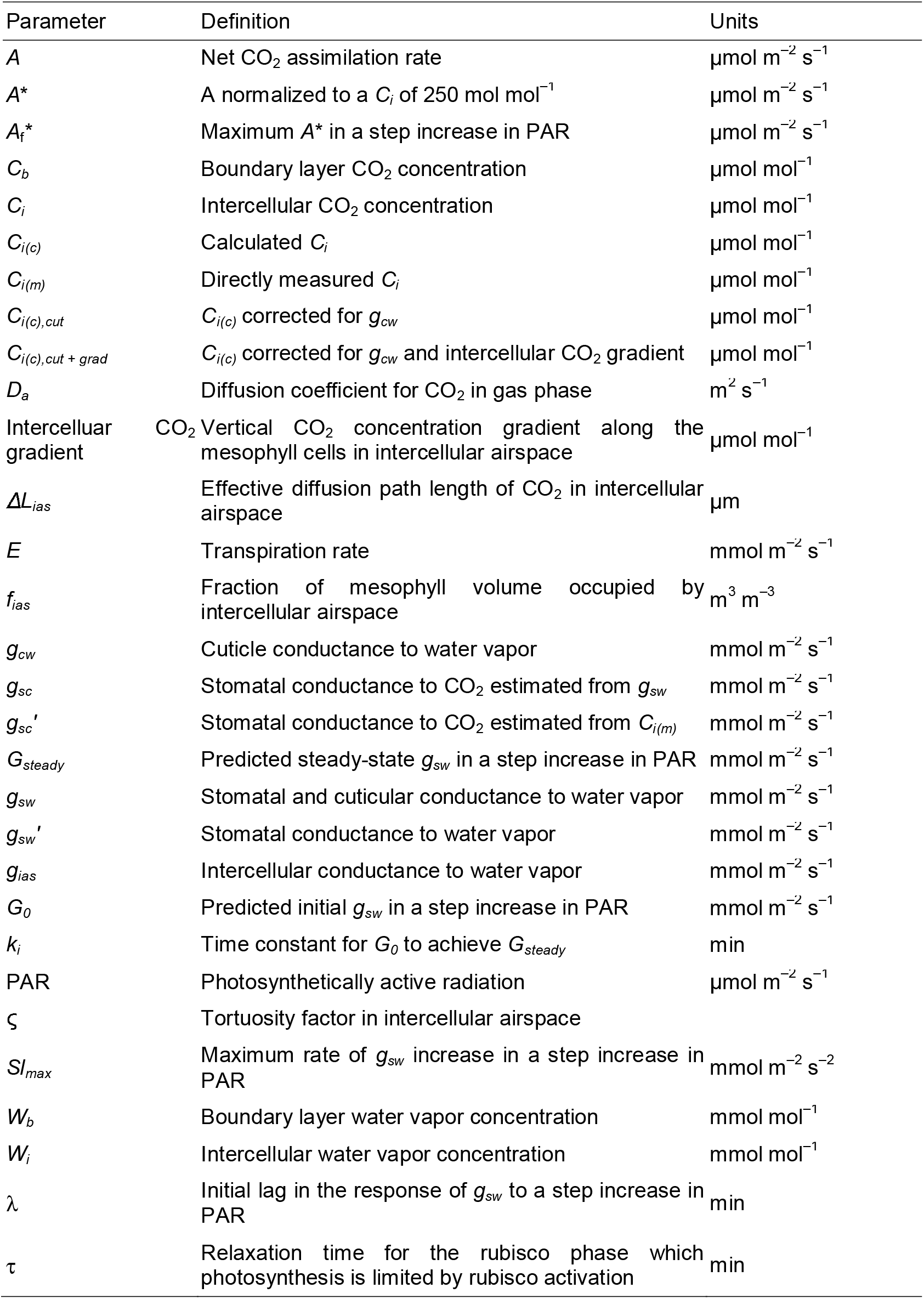
A summary of parameters referred to within the text.

### Stomatal kinetics

To get a sense of ‘rapidity’ of stomatal openings and increase of *g_sw_* in the leaves used here, we estimated kinetic parameters for the temporal response of *g_sw_* according to a sigmoidal model (McAusland et al., 2016; Vialet-Chabrand et al., 2017):

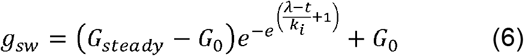

where *G_0_* and *G_steady_* are the *g_sw_* (mmol H_2_O m^−2^ s^−1^) at the time, *t* (min), for the beginning of light induction (*t* = 0) and at the steady state, respectively, *k_i_* (min) is the time constants for the increase of *g_sw_* and *λ* is the initial lag time (min). The parameter values were estimated with using a curve-fitting routine in Microsoft Excel provided elsewhere (Supplemental File S1 in Vialet-Chabrand et al., 2017). The maximum slope of *g_sw_* increase (*SI_max_*, mmol m^−2^ s^−2^) was calculated based on the previously described parameters as:

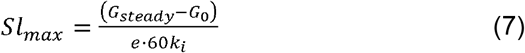

Our estimates only reflect the adaxial stomata while they usually represent both adaxial and abaxial stomata in amphi-stomatous leaves (e.g., McAusland et al., 2016).

### Rubisco activation kinetics

The activation of rubisco would limit photosynthesis within approximately the initial 10 min of the increase in PAR (Woodrow & Mott, 1989,1992). This phase can be found as a linear portion of a semilogarithmic plot of normalized photosynthesis to a constant *C_i_*(*A*\*) versus time (t) (Woodrow & Mott, 1989), expressed as:

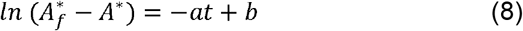

where *a* and *b* are the slope and intercept of the regression for this linear portion, 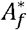 is the *A*\* in the final steady state. The relaxation time for the rubisco phase (*τ*) is then defined from the slope *a* as:

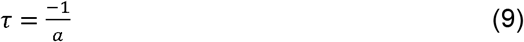

In the induction experiments from dark to saturating light, we estimated *τ* with using *C_i(c)_* or *C_i(m)_*, according to Woodrow and Mott (1989). *A* was normalized to a *C_i_* of 250 μmol mol^−1^ by assuming a linear relationship between *A* and *C_i_*, that passed through the CO_2_ compensation point of 50 μmol mol^−1^. The maximum *A*\* was used as 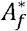 during each induction. *a* was determined by linear regression of the data for 3 min (i.e., 18 points).

### Stomatal density

Nail polish was spread at three points on each upper and lower side of ten leaves. The dried polish was striped with clear tape and attached to a glass slide (a replica of the epidermis). Microphotographs (0.60 mm^2^) of the replicates were taken with a digital camera (AxioCam HRc; Carl Zeiss, Göttingen, Germany) attached to a microscope (Axioskop 2 Mot Plus; Carl Zeiss, Göttingen, Germany). Stomata were counted in each replicate and averaged to calculate stomatal density for each side of the leaves.

### Intercellular CO_2_ conductance and gradients

Vertical gradients of CO_2_ exist in a leaf because CO_2_ has to further diffuse through the mesophyll airspace once inside the stomata (Parkhurst et al. 1988). In our measurement system, *C_i_* gets lower towards abaxial side because CO_2_ is only supplied from the adaxial side and more CO_2_ is consumed as it diffuses deeper into the leaf (Boyer and Kawamitsu, 2011). In one-dimensional model, the CO_2_ gradient is expressed as:

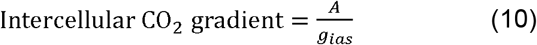

where *g_ias_* is the intercellular conductance of CO_2_ (mol CO_2_ m^−2^ s^−1^). We determined *g_ias_* anatomically from the fraction of mesophyll volume occupied by intercellular airspace (*f_ias_*) and the effective diffusion path length (*ΔL_ias_*) (Syvertsen et al., 1995) as:

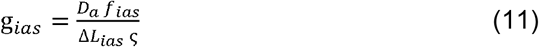

where *ζ* is the tortuosity factor and *D_a_* is the diffusion coefficient for CO_2_ in the gas phase (1.51·10^−5^ m^2^ s^−1^). We used mesophyll thickness as *ΔL_ias_* and a constant *ζ* of 1.57 (Tosens et al., 2012). The *f_ias_* and the mesophyll thickness between the two epidermal layers were measured from microscopic observations of leaf sections with the same equipment described above.

### Statistics

The number of replications and the statistical tests, if applicable, are presented in figure legends for each experiment. The results are given as means with SDs unless otherwise indicated.

## Results

### Leaf anatomy and intercellular conductance to CO_2_

Sunflower leaves had more stomata than tobacco leaves while their stomatal ratios were similar (Table 2). For adaxial sides on which gas exchange was measured, the mean stomatal density was 49 ± 4 mm^−2^ in tobacco and 118 ± 9 mm^−2^ in sunflower, respectively. In both species, adaxial side had fewer stomata than abaxial side with the mean stomatal ratio (adaxial/abaxial) of 0.6-0.7. Intercellular conductance of CO_2_ (*g_ias_*) was determined from anatomical properties of leaf sections (Table 2). Sunflower leaves trended toward having a smaller fraction of airspace (0.27 ± 0.01) than tobacco leaves (0.35 ± 0.05) whereas both leaves had a comparable mesophyll thickness of around 200 μm. There were no significant differences in the mean *g_ias_* between tobacco (617 ± 87 mmol CO_2_ m^−2^ s^−1^) and sunflower (538 ± 135 mmol CO_2_ m^−2^ s^−1^).

**Table 2.** Stomatal density and anatomical properties of tobacco and sunflower leaves.

|  | Tobacco | Sunflower | *P* |
| --- | --- | --- | --- |
| Stomatal density (mm<sup>-2</sup>, n=10) |  |  |  |
| Adaxial | 49±4 | 118±9 | <0.0001 |
| Abaxial | 76±16 | 186±20 | <0.0001 |
| Ratio (adaxial/abaxial) | 0.67±0.15 | 0.64±0.08 | 0.55 |
| Anatomical properties (n=3) |  |  |  |
| *f<sub>ias</sub>* | 0.35±0.05 | 0.27±0.01 | 0.06 |
| Mesophyll thickness (µm) | 225±53 | 199±3 | 0.49 |
| *g<sub>ias</sub>* (mmol m<sup>-2</sup> s<sup>-1</sup>) | 617±70 | 538±35 | 0.18 |

The means ± SDs are compared by using either Student's or Welch's *t* test.

### Light induction from dark to saturating irradiance

When dark-adapted leaves were clamped with the chamber under saturating irradiance (1200 PAR), both assimilation and conductance increased slowly (Fig. 1a,b). Meanwhile, CO_2_ concentration in the closed loop was rapidly consumed by photosynthesis until reaching an equilibrium with the airspace inside the leaf (*C_i(m)_* in Fig. 1c). Until then, the *C_i(m)_* was higher than the actual *C_i_* and the *g_sc_’* derived from the *C_i(m)_* was incorrect (shown with grey background in Fig. 1). The equilibrium state was indicated by the parallel changes in the *g_sw_* and *g_sc_’* as both changes should commonly reflect the stomatal behavior (inset in conductance of Fig. 1b). The equilibrium was generally expected to occur within 5 min after clamping on the leaf in both species. The *C_i(c)_* was significantly larger than the *C_i(m)_*, and the difference became largest when the *C_i(m)_* was around the minimum (Fig. 1c). The *C_i(m)_* decreased to 96 ± 15 μmol mol^−1^ in tobacco and 76 ± 11 μmol mol^−1^ in sunflower, whereas the minimum *C_i(c)_* was 137 ± 19 μmol mol^−1^ and 150 ± 10 μmol mol^−1^, respectively. Therefore, rubisco underwent much lower *C_i_* than was predicted from the calculation. Because rubisco activity can be inferred from *A* relative to *C_i_* within the early induction phase, these differences in the minimum *C_i_* would predict a different activation status of rubisco. Based on the *C_i(m)_* or *C_i(c)_*, we estimated the relaxation time (*τ*) primarily limited by the rate of rubisco activation as illustrated in Fig. S3. The *τ* estimated from the *C_i(c)_* (5.0 ± 2.7 min and 7.1 ± 2.8 min in tobacco and sunflower, respectively) were significantly longer than those estimated from the *C_i(m)_* (2.0 ± 1.0 min and 3.3 ± 2.0 min, respectively) (Fig. 2). The difference between the *C_i(c)_* and *C_i(m)_* diminished as the conductance increased (Fig. 1c).

**Fig. 1.**
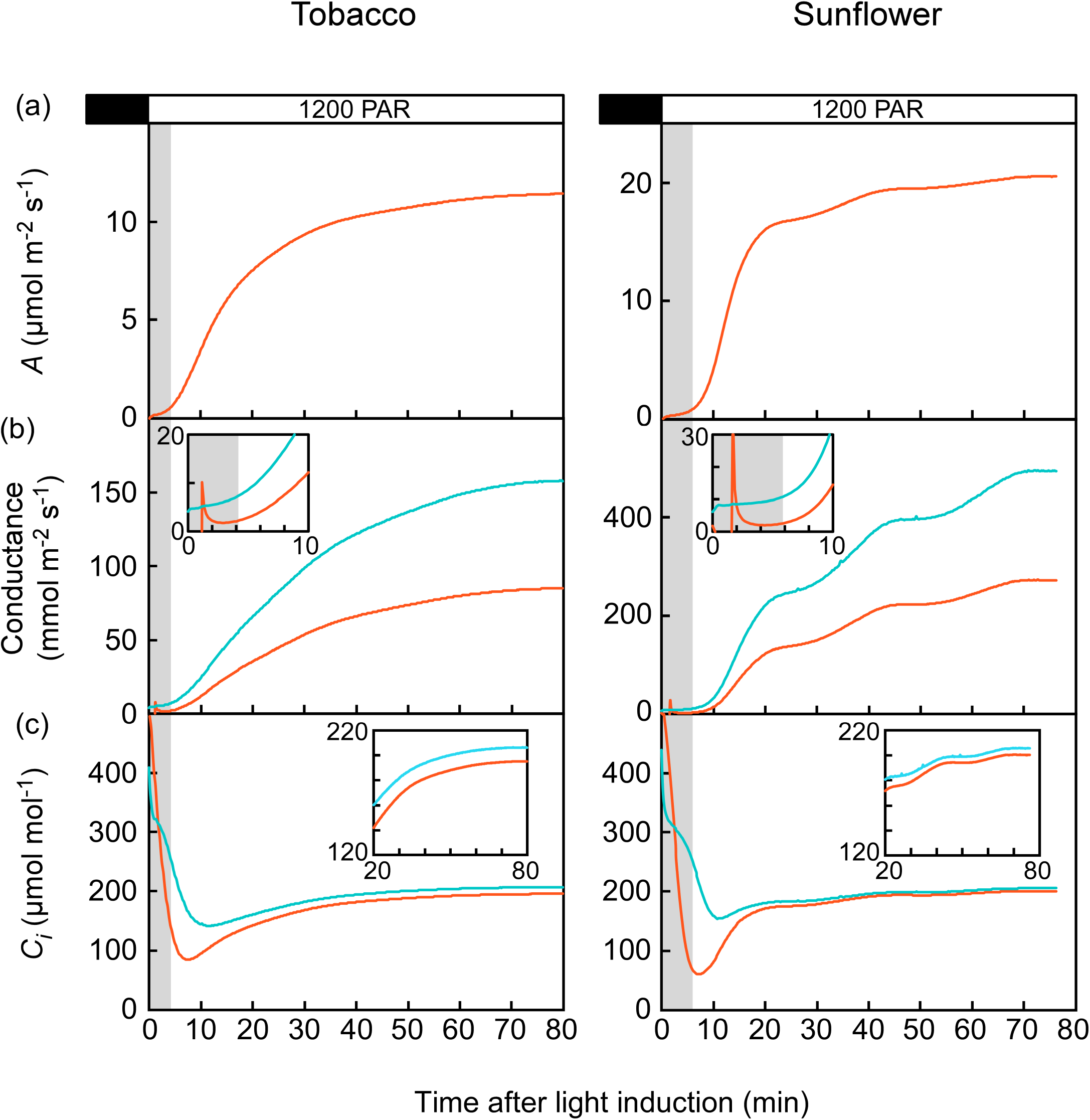
Changes in the standard gas exchange parameters during the light induction of photosynthesis from dark. Assimilation rate (*A*) (a), stomatal conductances to water vapor (*g_sw_*, blue line) and CO_2_ (*g_sc_*’, orange line) (b), and calculated intercellular CO_2_ concentration *(C_i(c)_*, blue line) and measured intercellular CO_2_ concentration *(C_i(m)_*, orange line) (c) after clamping a dark-adapted leaf of tobacco (left) and sunflower (right) with the dual gas-exchange chamber under saturating irradiance of 1200 μmol m^−2^ s^−1^ PAR. The *g_sw_* was derived from the standard calculation based on the water vapor flux, whereas the *g_sc_’* was derived from the direct measurement based on the CO_2_ flux (see Calculations). In (b), initial change (0-10 min) is zoomed in the inset. The x-axis ticks in the inset graph represent 2 min intervals. Pre-equilibrium time for the *C_i(m)_* was shown with grey background. Representative experiments from twelve and seven replications in tobacco and sunflower, respectively.

### Light induction from shade to saturating irradiances

In light induction experiments, leaves are often pre-acclimated to shade, rather than to the darkness. We also looked at the induction from shade (50 PAR) (Fig. 3). While the induction from dark to shade similarly resulted in a large discrepancy between the *C_i(c)_* and *C_i(m)_* as dark to saturating light, the subsequent induction from shade to saturating irradiance caused less discrepancies (Fig. 3c). However, tobacco *C_i(c)_* and *C_i(m)_* remained different at saturating light (insets of Figs. 1c & 2c). A large difference between the minimum *C_i(c)_* and *C_i(m)_* was also evident in the induction from the dark to 200 PAR (data not shown). Therefore, the calculations primarily deviated from the direct measurements only in the induction from the darkness. For those dark-adapted leaves, the initial *g_sw_* right after the induction was as small as 9.4 ± 6.6 mmol H_2_O m^−2^ s^−1^ and 9.4 ± 6.8 mmol H_2_O m^−2^ s^−1^ for *g_sw_* in tobacco and sunflower, respectively. On the other hand, when the photosynthesis reached a steady-state in the shade, *g_sw_* increased to as much as 61 ± 20 mmol m^−2^ s^−1^ in tobacco, and 190 ± 56 mmol m^−2^ s^−1^ in sunflower. These results are consistent with the previous direct measurements showing the over-estimation of *C_i(c)_* mostly when *g_sw_* is small (Tominaga & Kawamitsu, 2015b, Boyer, 2015a; Tominaga et al., 2018), and these initial *g_sw_* in the shade-acclimated leaves must have been sufficiently large to avoid over-estimations.

**Fig. 2.**
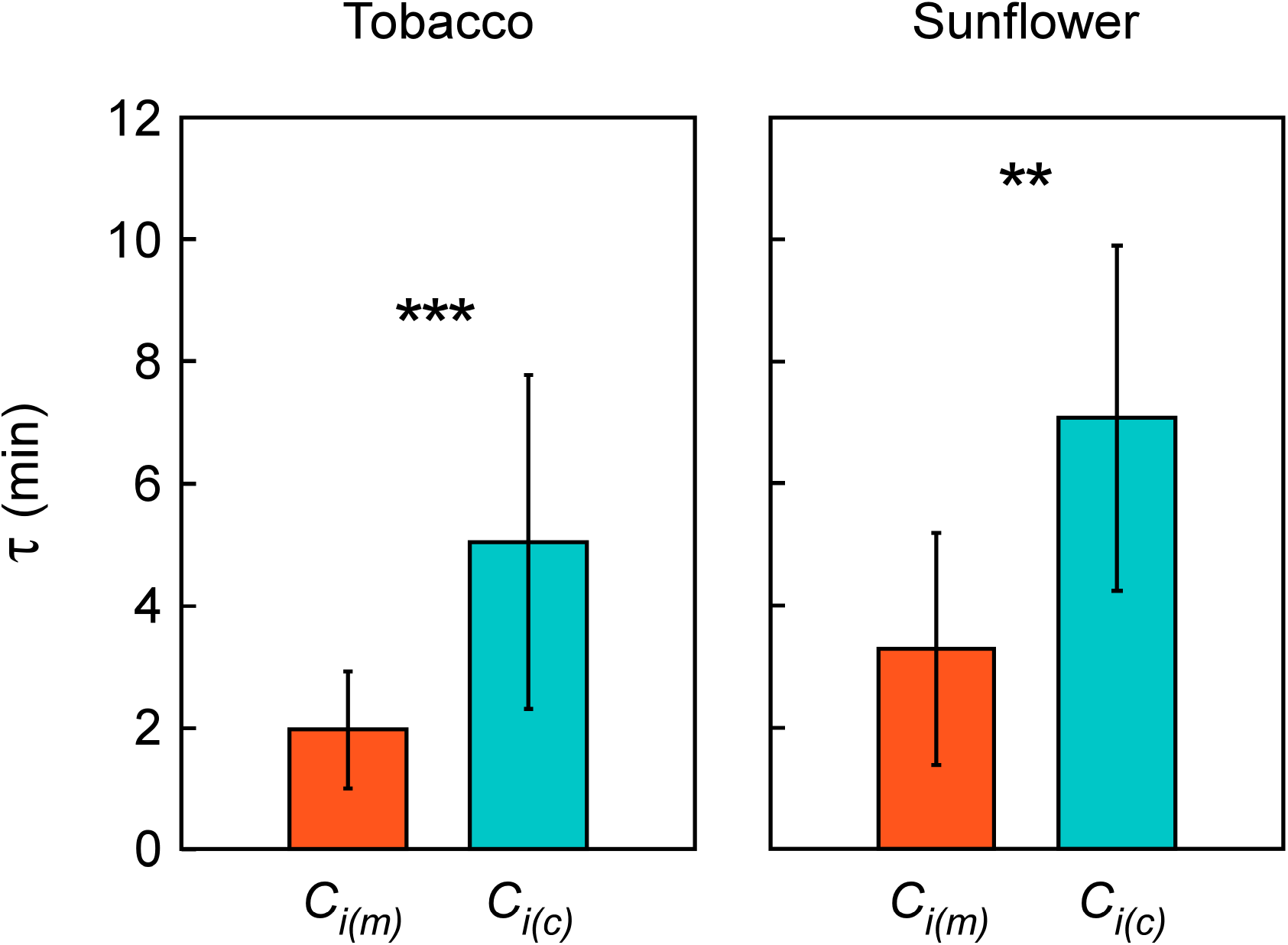
Relaxation time (*τ*) for the rubisco phase at the minimum *C_i(m)_* (orange) or *C_i(c)_* (blue). Means ± SDs (n=12 and 7 in tobacco and sunflower, respectively) were compared by using t test (***: P<0.001**: P<0.01).

### Kinetics of stomatal openings

Kinetic parameters for temporal response of *g_sw_* during the light induction were summarized in Table 3. In general, increase of *g_sw_* was sigmoidal in tobacco whereas it was multiple stepwise in sunflower (Figs. 1b & 3b). As a result, the smooth sigmoidal function (Eq. 6) was less fit in sunflower showing the higher normalized root-mean-square error (NRMSE). Time constant of stomatal openings (*k_i_*) was shorter in the induction from dark to shade (0 to 50 PAR), suggesting that stomata reached steady-state more quickly with partial openings. On the other hand, the lower *SI_max_* indicated the slower increase of *g_sw_* in the shade. In the induction from dark to saturating light (0 to 1200 PAR), average *SI_max_* was higher in sunflower (0.139 mmol m^−2^ s^−2^) than in tobacco (0.047 mmol m^−2^ s^−2^) despite the similar *k* (approx. 20 min) because of the larger steady-state *g_sw_* with the higher stomatal density (Table 2). These *SI_max_* values were within or close to the high end of reported values for C_3_ forbs displaying the similar growth form and guard cell type (McAusland et al., 2016). Hence, it follows that over-estimations of *C_i(c)_* in the above experiments occurred in leaves with moderate to rapid *g_sw_* responses. Shade acclimation (50 to 1200 PAR) neither shortened the *k_i_* nor increased the *SI_max_* (Table 3).

**Fig. 3.**
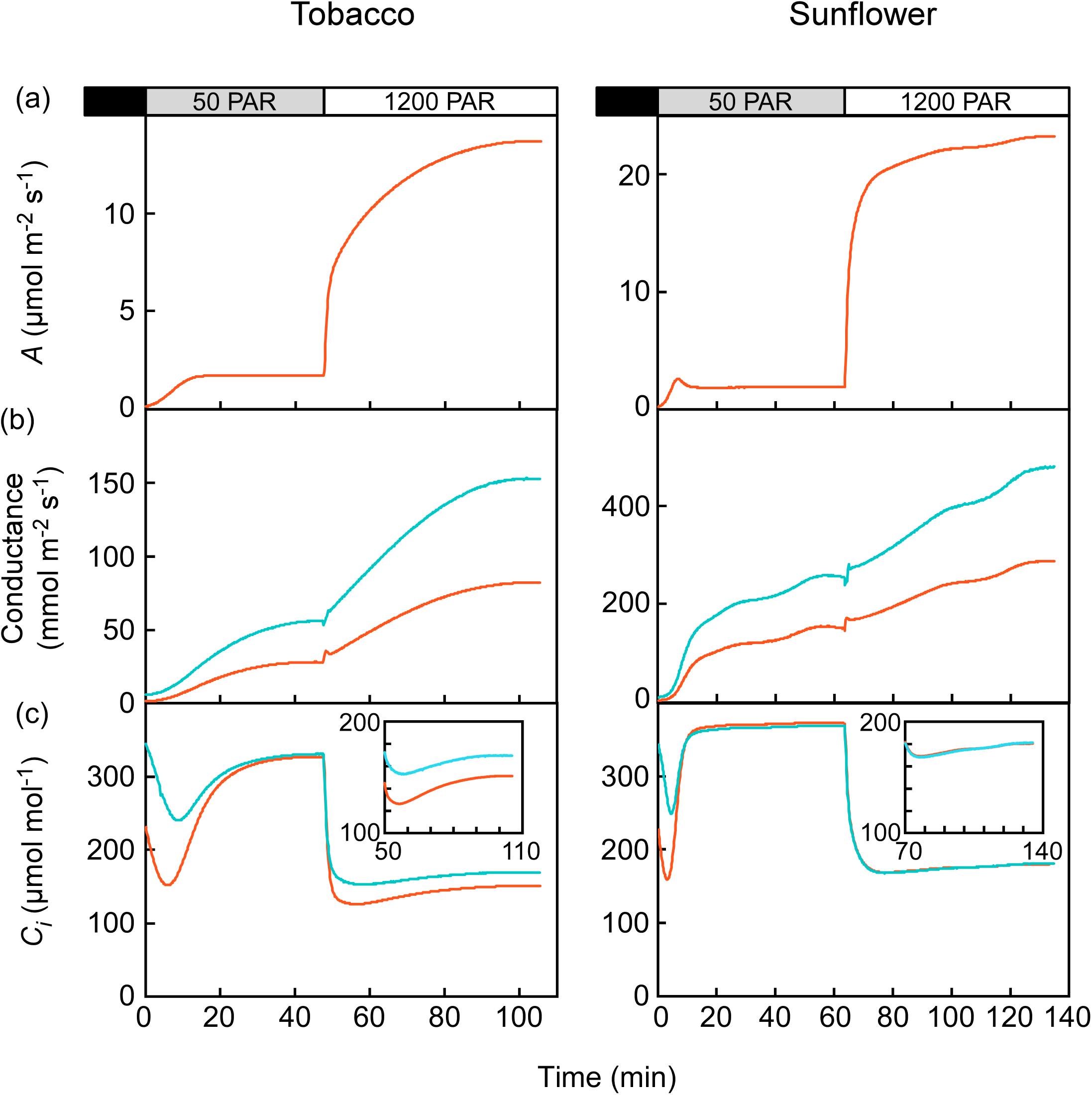
Light induction experiments from dark to a shade (50 μmol m^−2^ s^−1^ PAR) and to a saturating irradiance (1200 μmol m^−2^ s^−1^ PAR). *A* (a), *g_sw_* (blue line) and *g_sc_’* (orange line) (b), and *C_i(c)_* (blue line) and *C_i(m)_* (orange line) (c). Measurements prior to the equilibrium were removed. Representative experiments from six and four replications in tobacco and sunflower, respectively.

**Table 3.** Parameters of the dynamic model of *g_sw_* increase during light induction experiments in tobacco and sunflower.

| PAR<br>(µmol m<sup>-2</sup> s<sup>-1</sup>) | Species | n | NRMSE | *k<sub>i</sub>*<br>(min) | λ<br>(min) | *S<sub>I</sub>*<sub>max</sub><br>(mmol m<sup>-2</sup> s<sup>-2</sup>) | *G<sub>0</sub>*<br>(mmol m<sup>-2</sup> s<sup>-1</sup>) | *G<sub>steady</sub>*<br>(mmol m<sup>-2</sup> s<sup>-1</sup>) |
| --- | --- | --- | --- | --- | --- | --- | --- | --- |
| 0 to 50 |  |  | ** |  |  |  |  | *** |
|  | Tobacco | 6 | 0.018 ± 0.006<sup>bc</sup> | 11.3 ± 1.2<sup>b</sup> | 1.9 ± 1.0<sup>a</sup> | 0.034 ± 0.004<sup>b</sup> | 3.0 ± 1.8<sup>b</sup> | 64.4 ± 8.5<sup>b</sup> |
|  | Sunflower | 4 | 0.057 ± 0.010<sup>A</sup> | 15.5 ± 3.8<sup>B</sup> | 2.1 ± 1.7<sup>A</sup> | 0.092 ± 0.029<sup>AB</sup> | 7.3 ± 7.3<sup>B</sup> | 188.5 ± 26.3<sup>C</sup> |
| 0 to 200 |  |  |  |  | * | *** |  | *** |
|  | Tobacco | 9 | 0.032 ± 0.002<sup>ab</sup> | 17.8 ± 1.1<sup>a</sup> | 0.3 ± 0.2<sup>ab</sup> | 0.050 ± 0.004<sup>a</sup> | 6.6 ± 6.4<sup>b</sup> | 149.7 ± 7.7<sup>a</sup> |
|  | Sunflower | 8 | 0.057 ± 0.007<sup>A</sup> | 18.0 ± 1.7<sup>AB</sup> | 5.4 ± 2.1<sup>A</sup> | 0.134 ± 0.013<sup>A</sup> | 0.0 ± 0.0<sup>B</sup> | 369.6 ± 16.6<sup>B</sup> |
| 0 to 1200 |  |  |  |  |  | *** |  | *** |
|  | Tobacco | 12 | 0.035 ± 0.004<sup>a</sup> | 20.0 ± 1.1<sup>a</sup> | 0.0 ± 0.0<sup>b</sup> | 0.047 ± 0.003<sup>ab</sup> | 12.8 ± 11.4<sup>b</sup> | 165.4 ± 9.9<sup>a</sup> |
|  | Sunflower | 7 | 0.048 ± 0.007<sup>A</sup> | 18.8 ± 2.2<sup>AB</sup> | 1.9 ± 1.3<sup>A</sup> | 0.139 ± 0.013<sup>A</sup> | 7.1 ± 2.9<sup>B</sup> | 410.7 ± 28.9<sup>AB</sup> |
| 50 to 1200 |  |  | ** | ** |  | ** | ** | *** |
|  | Tobacco | 9 | 0.012 ± 0.002<sup>c</sup> | 19.1 ± 1.2<sup>a</sup> | 0.8 ± 0.5<sup>ab</sup> | 0.041 ± 0.004<sup>ab</sup> | 54.3 ± 16.7<sup>a</sup> | 178.7 ± 11.4<sup>a</sup> |
|  | Sunflower | 4 | 0.032 ± 0.007<sup>A</sup> | 27.1 ± 1.3<sup>A</sup> | 0.0 ± 0.0<sup>A</sup> | 0.066 ± 0.002<sup>B</sup> | 190.9 ± 24.7<sup>A</sup> | 484.1 ± 34.9<sup>A</sup> |

Means ± SEs. Different letters in each species indicate significant (*P* < 0.05) difference among the irradiance according to Tukey-Kramer test, whereas star(s) in each experiments indicate significant (*P* < 0.05) difference between the species according to *t* test (\*\*\*: *P* < 0.001, \*\*: *P* < 0.01, \*: *P* < 0.05). Note that McAusland et al. (2016) reported 6.9-23.4 min and 0.04-0.13 mmol m<sup>-2</sup> s<sup>-1</sup> for the range of *k<sub>i</sub>* and *S<sub>I</sub>*<sub>max</sub>, respectively, in the light induction from 100 to 1000 PAR for several C<sub>3</sub> forbs.

### Estimation and correction of cuticle conductance

The standard calculations use the diffusivity ratio of water vapor to CO_2_, which is a constant value of 1.6 in air (Sharkey & Farquhar, 1982; LI-COR, 2019) while assuming that stomata are the dominant path for both gasses (Eq. (2)). Note that Massman (1998) reported uncertainty (maximum deviations after removing outliers) of ± 7% and ± 5% for the respective diffusivities, indicating ± 12% uncertainty (± 0.19) in the mean ratio of 1.58. We tested this assumption that *g_sw_* = 1.6*g_sc_*’ by comparing the *g_sw_* with the *g_sc_’* during the initial light induction from the dark (*g_sc_*’<20 mmol CO_2_ m^−2^ s^−1^) (Fig. 4). The *g_sw_* was linearly related to the *g_sc_’*, indicating a tight stomatal regulation on the *g_sw_* in this early induction phase. However, the *g_sw_* was consistently larger than the assumption of *g_sw_* = 1.6*g_sc_*’ (broken lines in Fig. 4). Given little CO_2_ transport through the cuticle (Boyer, 2015a), a zero *g_sc_’* would be obtained with complete stomatal closure. Thus, the *g_sw_* with complete stomatal closure can be estimated by extrapolating the regression line to the intersection with the y-axis. This intersection is consistently above the origin (Fig. 4), indicating that water transport continues even if stomata close completely. We considered this residual *g_sw_* as cuticle conductance for water vapor (*g_cw_*).

**Fig. 4.**
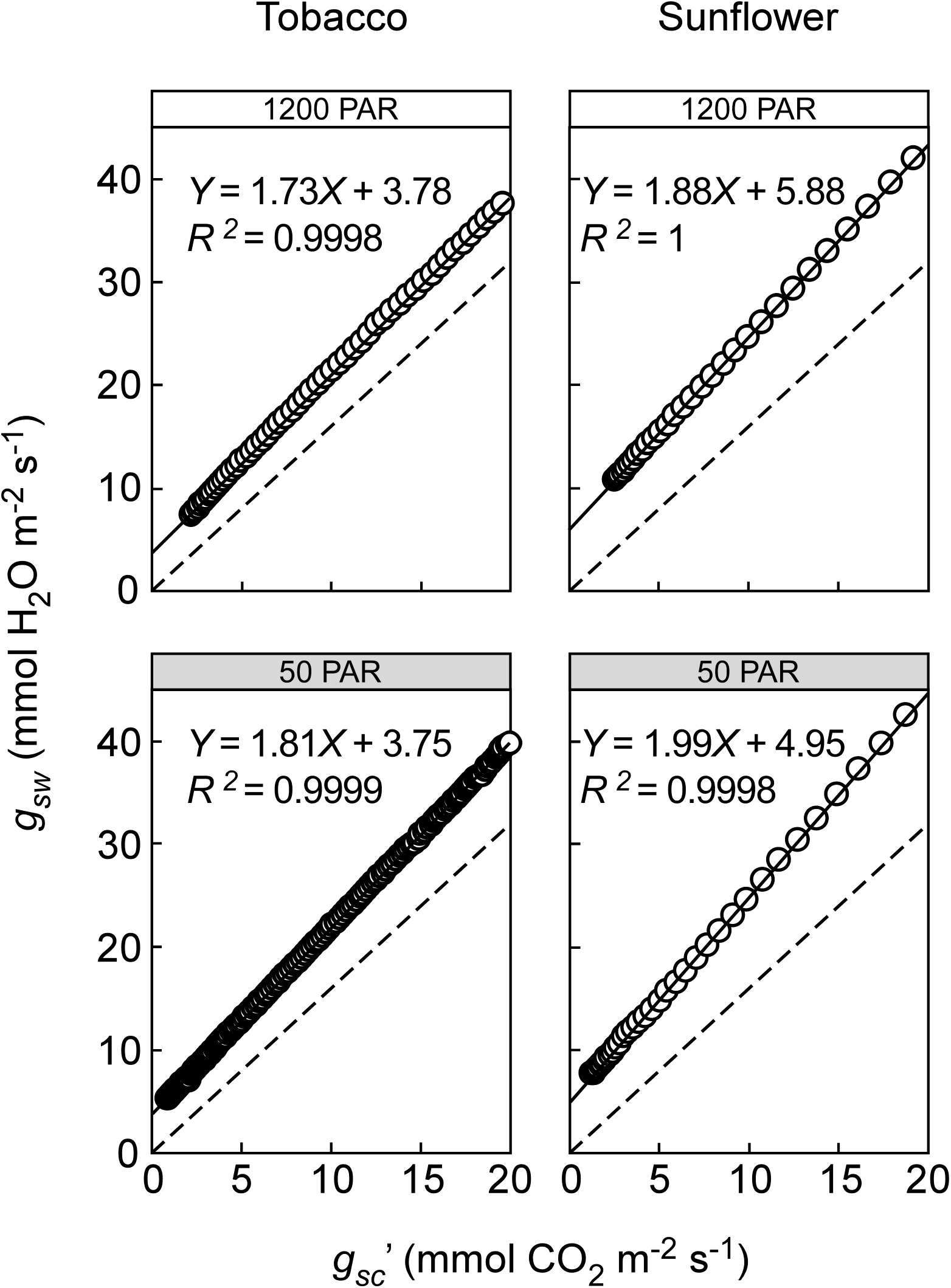
Relationships between *g_sw_* and *g_sc’_* during initial phase (*g_sc_’* < 20 mmol CO_2_ m^−2^ s^−1^) of the light induction from the dark (< initial 10 min). Data for Figs. 1 and 3 are shown. Pre-equilibrium section was excluded as described in the text. Broken lines indicate the relationship for the diffusivity ratio of water vapor to CO_2_ of 1.6 (*g_sw_* = 1.6 *g_sc_’*). Cuticle conductance to water (*g_cw_*) was extrapolated to be y-intercept of the regression line.

To see the impact of cuticle conductance, we corrected *C_i(c)_* by calculating the strict sense of stomatal conductance to water vapor (*g_sw_*’), as described in Eq.(4), and substituting it for the *g_sw_*. The initial extrapolated *g_sw_’* right after the induction from dark was 6.8 ± 6.5 mmol H_2_O m^−2^ s^−1^ and 6.1 ± 6.4 mmol H_2_O m^−2^ s^−1^ in tobacco and sunflower, respectively, that are 63 ± 21% and 56 ± 22% of the initial *g_sw_*. The corrected *C_i(c)_* (*C_i(c),cut_*) followed the *C_i(m)_* more closely than the *C_i(c)_* and the discrepancy became much smaller (Fig. 5a,b), suggesting that the discrepancy in the early induction was mostly attributable to the contribution of cuticular water loss. In contrast, the *g_cw_* marginally corrected the *C_i(c)_* as the *g_sw_* increased, and there remained the discrepancy even when the *g_sw_* was maximum, especially in tobacco (insets of Fig. 5a,b).

**Fig. 5.**
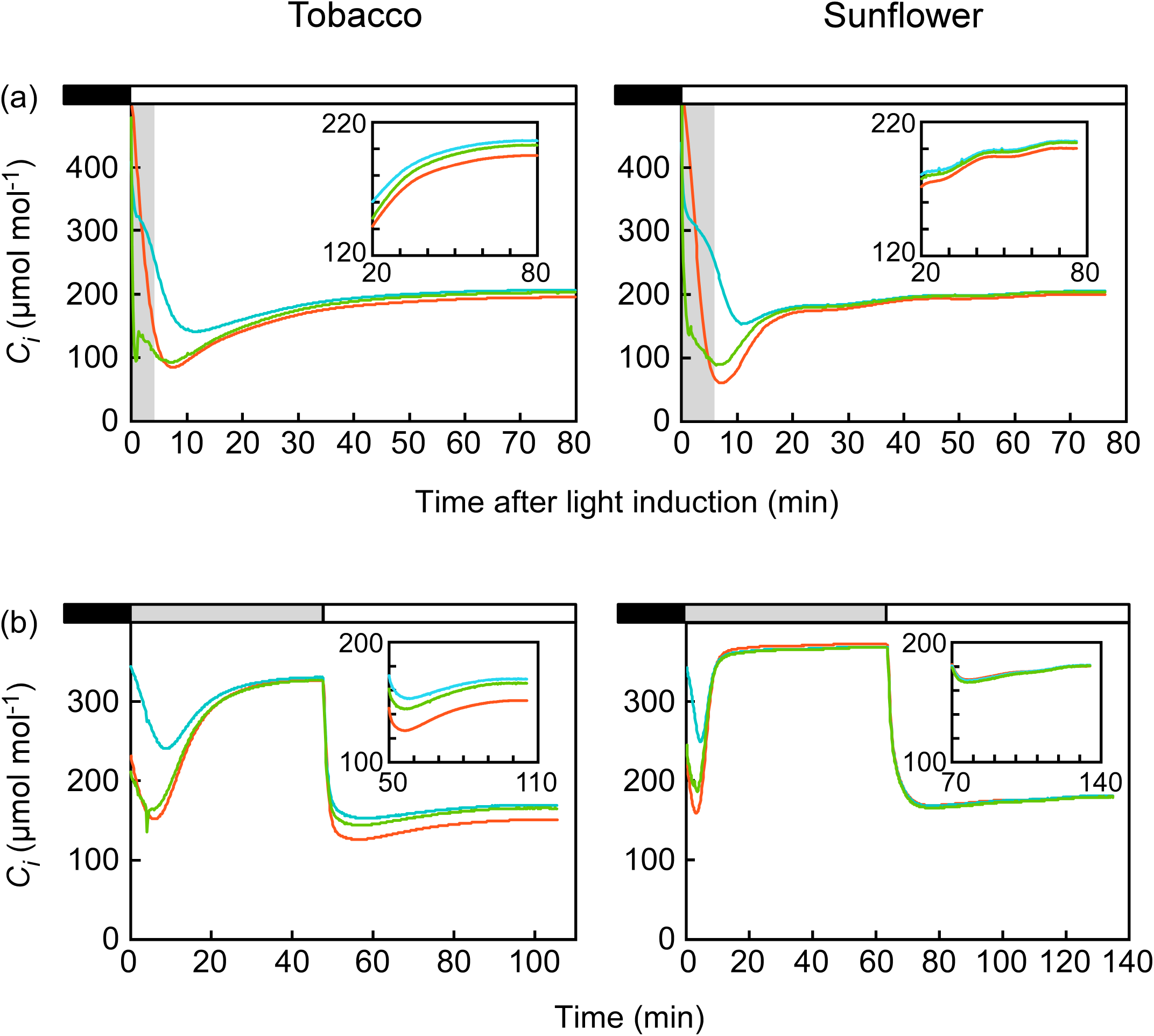
Correction for cuticle conductance in light induction experiments. Data for Figs. 1 and 3 are shown in (a) and (b), respectively. The *C_i(c)_* was corrected (*C_i(c),cut_*,green line) by estimating the strict sense of stomatal conductance to water vapor (*g_sw_’*) with the estimated *g_cw_* at each measurement, according to Eq. (4).

The *g_cw_* did not show a particular trend in response to the irradiance, with the mean across all irradiances of 2.65 ± 1.10 mmol m^−2^ s^−1^ in tobacco and 3.27 ± 1.93 mmol m^−2^ s^−1^ in sunflower, respectively (Fig. 6a). Because of the strict linearity of the regressions (Fig. 4), the standard error of each *g_cw_* estimate (i.e., y-intercept) was less than 0.1 mmol m^−2^ s^−1^ in both species, that is much smaller than the variations among leaves (Fig. 6a). The slope of the regression was mostly within or slightly above the uncertainty interval of 1.58, except for the inductions in shade (Fig. 6b). The slower photosynthesis in the shade (Table S1) would delay the equilibrium in the closed system, and perhaps affected the slope. The slope value was greater in sunflower than in tobacco across the irradiances (Fig. 6b).

**Fig. 6.**
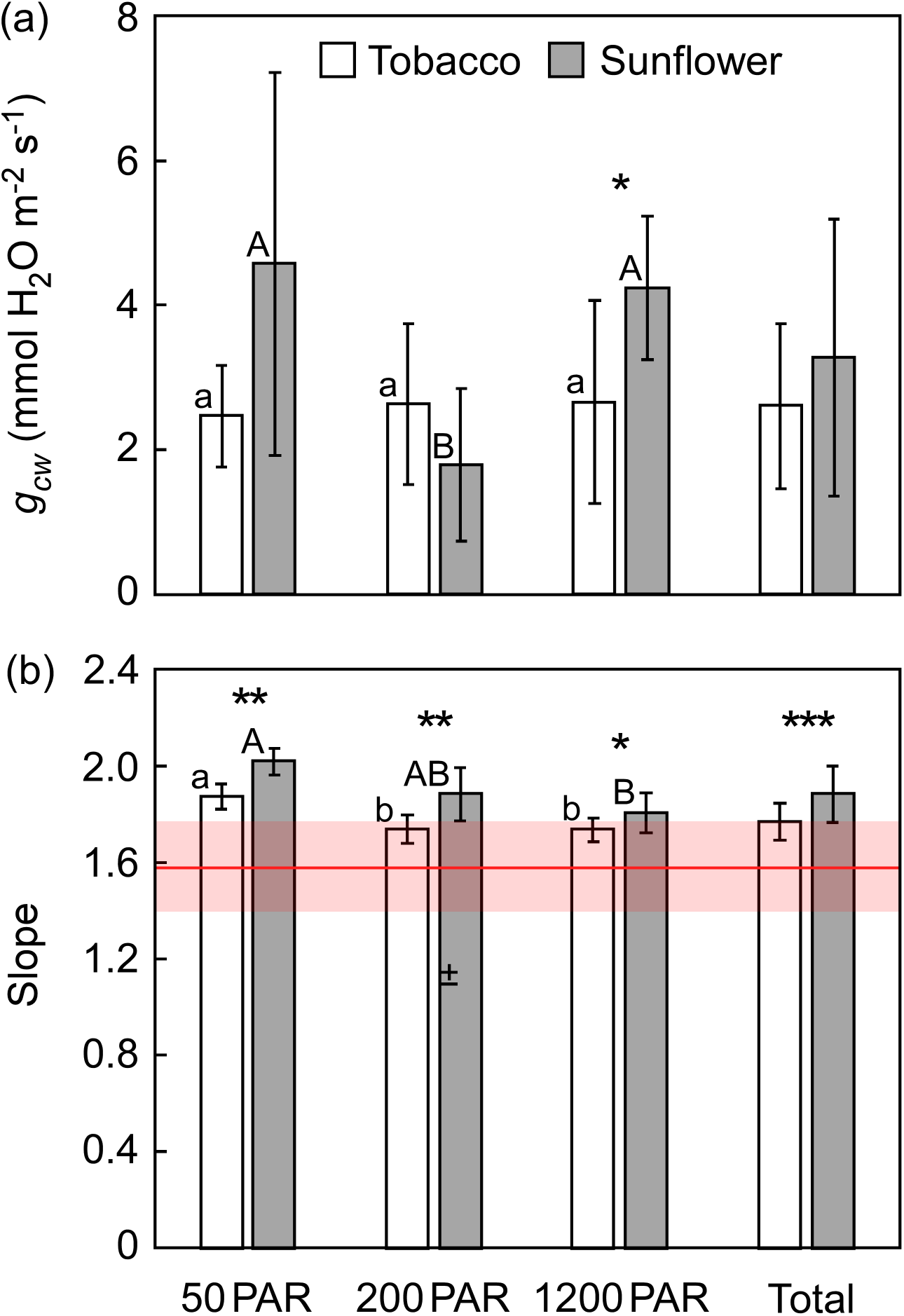
Summary of *g_cw_* (a) and slope (b) estimated from the regression line for the *g_sw_* versus *g_sc_* relationship at various irradiances. Means ± SDs [n=6 and 4 (50 PAR), n=9 and 8 (200 PAR), and n= 12 and 7 (1200 PAR), in tobacco and sunflower, respectively]. In (b), the modeled diffusivity ratio of water vapor to CO_2_ in air (1.58) and its uncertainty intervals (± 12%) (Massman, 1998) were shown with a horizontal red line and pale red color, respectively. Different letters in each species indicate significant (P < 0.05) difference among the irradiance according to Tukey-Kramer test, whereas star(s) in each experiment indicate significant (P < 0.05) difference between the species according to t test (***: P<0.001**: P<0.01, *: P<0.05).

### Effect of intercellular CO_2_ gradients

Because intercellular conductance is finite, vertical gradients of *C_i_* must be present in a leaf. In our measurement system, the *C_i(m)_* should be the lowest *C_i_* as the CO_2_ is only supplied from the opposite side, and the CO_2_ gradients would be attributable to the residual difference of *C_i(c)cut_* from *C_i(m)_*. Assuming that the *C_i(c)_* represents the value just inside the stomatal pore or ‘stomatal cavity’ on the adaxial stomata (Sharkey et al., 1982; Parkhurst et al; 1988; Buckley et al., 2017), we subtracted the gradients estimated with the mean *g_ias_* (Table 2) according to Eq. (10) from the *C_i(c),cut_*. The gradient reduced the *C_i(c)_* when the *g_sw_* was relatively large (*C_i(c),cut + grad_* in Fig. 7a,b). The gradients slightly overcorrected the *C_i(c)_* in tobacco (insets of Fig. 7a,b), and these mismatches were adjustable by approximately doubling the *g_ias_* (data not shown). In sunflower the *C_i(c)_* was already close to or even lower than the *C_i(m)_* by several-ppm when the stomata were open (Figs. 1c & 2c). Then, the gradients greatly overcorrected the *C_i(c)_* (Fig. 7a,b), and the *g_ias_* had to be unrealistically large—more than 10X larger than the anatomical *g_ias_* (>5000 mmol m^−2^ s^−1^)—in order to match the *C_i(c),cut+grad_* with the *C_i(m)_*. Therefore, the over-correction would rather indicate that *C_i(c)_* was indicated deeper inside the leaf than the adaxial stomatal cavity and/or that *C_i(c)_* was underestimated.

**Fig. 7.**
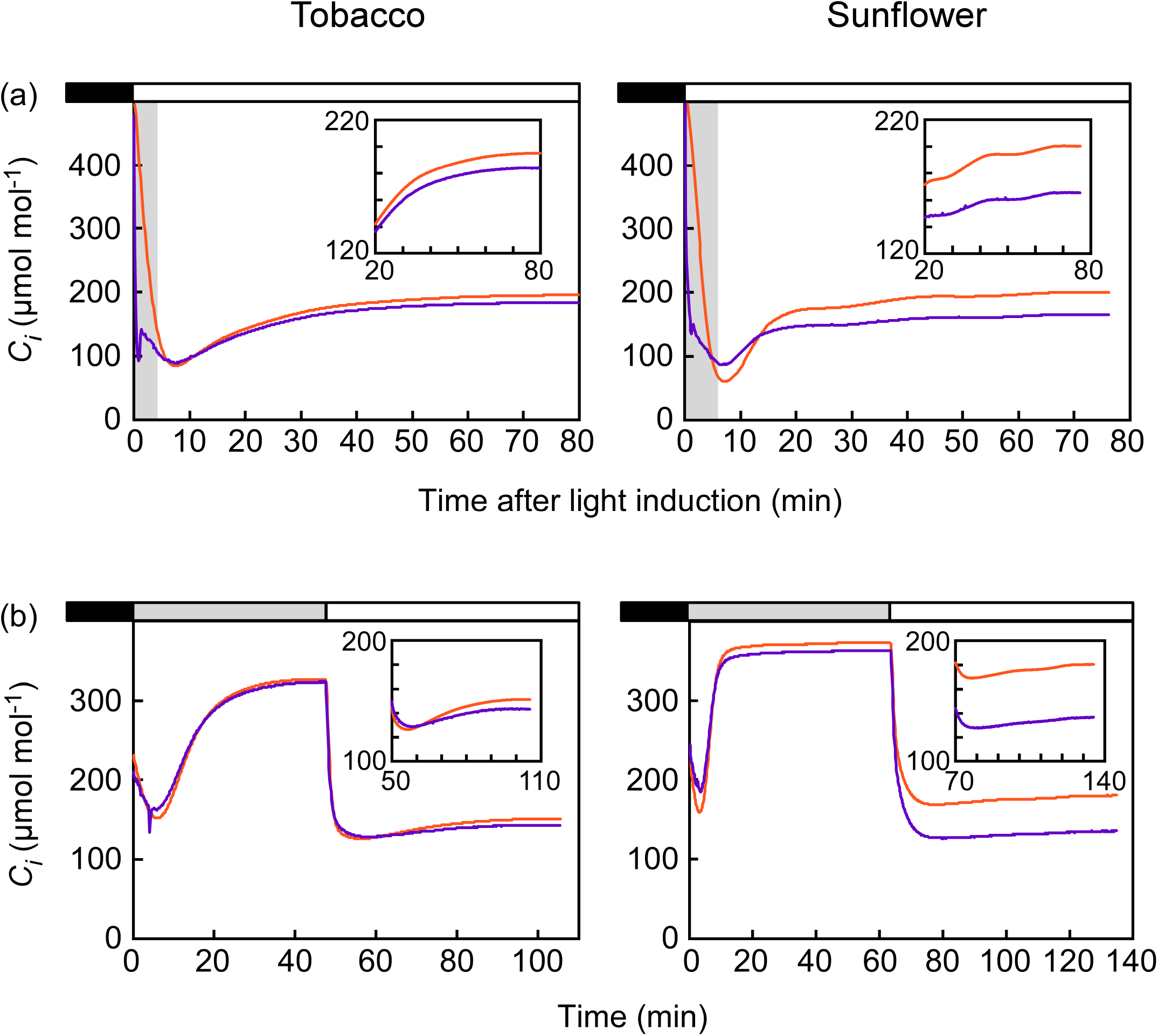
Effect of intercellular CO_2_ gradients on the calculation in light induction experiments. Data for Figs. 1 and 3 are shown in (a) and (b), respectively. The *C_i(c)_* was corrected for CO_2_ gradients after the correction for cuticle effect (*C_i(c),cut + grad_*, purple line) by subtracting the CO_2_ gradient estimated with the mean *g_ias_* (Table 1) from the *C_i(c),cut_*. Comparisons are *C_i(m)_* (orange line).

## Discussion

In this study, the intercellular CO_2_ concentration inside leaves was directly measured as well as calculated from the standard gas exchange measurement. A commercially available open system was minimally modified to include the direct system, and simultaneous measurements on the same leaf area allowed a close comparison. The standard calculations for the gas exchange measurements were validated under a square-wave light pattern that is often employed for induction studies. Over-estimations of *C_i_* were observed in the dark-adapted leaves whose stomata started out closed. This raises a potential problem for interpreting gas exchange kinetics during non-steady state photosynthesis because *C_i_* is an essential indicator of dynamic limitations (Kirshbaum & Pearcy, 1988; Pearcy et al., 1996). In our experiments, over-estimation of *C_i_* was profound most at around its minimum during the early induction phase (Fig. 1c). Within this timescale photosynthesis is primarily limited by the rate at which rubisco is converted from an inactive to an active form (Woodrow & Mott, 1989, 1992). We found that the over-estimation of the minimum *C_i_* delayed the relaxation period of the rubisco phase (Fig. 2). This suggests that rubisco activation status derived from *A* vs *C_i_* relationship is potentially underestimated while the rubisco actually experiences lower CO_2_ concentrations than the calculation indicates. In other words, one would be fooled as if rubisco is decarbamylated at the higher *C_i(c)_* than the actual *C_i_*. Our study demonstrated that the cuticular water loss was responsible for the calculation error and should be taken into account in the induction measurements.

Importantly, the *g_cw_* hardly affected the induction calculation when stomata were partially opened in the shade prior to the full induction (cf. *C_i(c)_* & *C_i(c),cut_* in Fig. 5b). In the induction from shade in *Alocasia macrorrhiza* (giant taro), Kirshbaum and Pearcy (1988) also realized that the *A/C_i_* slope (at the initial 1 min) declined when the initial *g_sw_* was suppressed under high CO_2_. While stomata would partially open in fairly low irradiance under the current atmospheric CO_2_ level (Valladares et al., 1997), the initial *g_sw_* varies greatly among species in both shade (McAusland et al., 2016; Deans et al., 2018) and darkness (Wachendorf & Küppers, 2017). The initial *g_sw_* is confined in and likely scaled to the maximum *g_sw_* among taxa in vascular plants (McAusland et al., 2016, Deans et al., 2019). Because greater stomatal conductance appears to evolve through the vascular plant lineages (Flanks & Beerling, 2009; McElwain et al., 2016), ferns and gymnosperms may have smaller initial *g_sw_* than angiosperms. Together, ‘sluggish’ stomatal openings could exacerbate the calculation by prolonging the cuticle effect. In general, graminoids with dumbbell-shaped guard cells exhibit much faster light response of *g_sw_* than the other vascular plants with elliptic guard cells (Franks & Farquhar, 2007; Vico et al., 2011; McAusland et al., 2016). In the elliptic guard cells, both stomatal opening and *g_sw_* increase are slower with lower maximum *g_sw_* (McAusland et al., 2016), and gymnosperm opens stomata more slowly than angiosperm (Vico et al., 2011). Besides genetic variations, water stress decreases the initial *g_sw_* (Allen & Pearcy, 2000; Wachendorf & Küppers, 2017) and so drought conditions should be prone to the error.

Cuticle conductance also varies among leaves in a species (Fig. 6a) and potentially more among species (Kerstiens, 1996; Schuster et al., 2017). Furthermore, intact cuticle conductance may change with leaf water status (Boyer et al., 1997; Burghardt & Riederer, 2003; Boyer, 2015b). Therefore, a correction seems problematic unless *g_cw_* for each intact leaf is known. Indeed, the average *g_cw_* could result in over-correction of *C_i_* in our experiments (data not shown). Although, in principle, the cuticle conductance can be easily included in the calculations for *C_i_* (*C_i(c),cut_*), accurate measurement of *g_cw_* is essential for a reliable correction.

### Estimation and correction of cuticle conductance

Due to technical difficulty in the measurement of cuticular transpiration on a stomatous leaf surface/cuticle (e.g., Šantrůček et al., 2004), cuticle conductance has been studied exclusively in hypostomatous species with using the astomasous leaf surface/cuticle (Kerstiens, 1996; Schuster et al., 2017). Kirshbaum and Pearcy (1988) first estimated the *g_cw_* on the stomatous leaf surface from the above induction measurements in giant taro. They determined the *g_cw_* that could match the two initial *A*/*C_i_* slopes observed with a low and high *g_sw_* in each leaf, and reported the mean of 2.7 ± 0.3 mmol m^−2^ s^−1^ (± SE, n=7). As they noted, however, the method assumes that the activation of rubisco is independent from the initial *g_sw_*, which would not hold (Mott & Woodrow, 1993). In the previous experiments using similar dual gas-exchange systems (Boyer 2015a, Tominaga & Kawamitsu, 2015b), *g_cw_* or cuticular transpiration was estimated from a deviation of the calculation from the direct measurement. Unlike the Kirshbaum and Pearcy method, these estimations can be made from a single measurement without assuming the rubisco activation status, thereby increasing the throughput. Nevertheless, these methods still assumed that *g_cw_* is solely responsible for their measurement discrepancies, and potentially neglected other factors, such as intercellular CO_2_ gradients. In contrast, our intercept method extrapolates *g_cw_* at virtually complete stomatal closure when the CO_2_ gradient becomes negligibly small with *A*≈0 (Fig. 4). Furthermore, the linear relationship allowed a simple extrapolation for a correction that improved the agreement between calculated and measured values (Fig. 5).

The regression slope was slightly larger than 1.58, the modelled diffusivities ratio of CO_2_ and water vapor in air (Massman, 1998), but mostly within the uncertainty interval (Fig. 6b). A potential explanation for the larger slope is a non-uniform stomatal opening that breaks the assumptions of lateral uniformity over the leaf surface: uniform stomatal aperture, leaf temperature and metabolic capacity, and in the leaf: uniform CO_2_ distribution (Terashima et al., 1988; Meyer & Genty, 1998; West et al., 2005). Because non-uniformity may be more obvious in heterobaric leaves in which bundle-sheath extensions laterally disconnects the airspace (Terashima, 1992), the larger slope in sunflower than tobacco (Fig. 5b) might be due to the greater heterobaricity (Morison & Lawson, 2007; also noticeable in our microscopic observation). It would be interesting to compare the thermal and fluorescence imaging over the leaf surface (e.g., West et al., 2005) during induction experiments among leaves with a wide range of heterobaricity.

### Intercellular CO_2_ gradients and unsaturation in the airspace

In tobacco, the intercellular CO_2_ gradient seems to explain the discrepancy between the *C_i(c)_* and *C_i(m)_* when stomata are open and *g_cw_* has little impact on the calculation (Fig. 7). The slight over-correction of *C_i(c)_* may be due to under-estimation of the anatomical *g_ias_*. However, the much larger over-correction of *C_i(c)_* in sunflower would indicate that the *C_i(c)_* was located deeper than the stomatal cavity (i.e., closer to the *C_i(m)_* than we expected). This suggests that internal evaporating surface where humidity is saturated (*W_i_*) occurs deeper than the stomatal cavity, and that *g_sw_* includes the diffusion in the intercellular airspace. Also, this notion could explain the discrepancy in the over-correction between the two species. Because closer arrangements of stomata should decrease the lateral diffusion distance of CO_2_ inside the leaf, the pathlength ratio of water vapor to CO_2_ per stoma may increase with increasing stomatal density. In sunflower leaves having the 2.4X as many stomata (Table 2), a larger pathlength ratio might cause the deeper *C_i_*.

It is also possible that the over-correction is manifested by under-estimated *C_i(c)_*. In calculating the *g_sw_* (Eq. 3), *W_i_*, is assumed to be saturated, and if not the *g_sw_* is more or less underestimated. We observed a progressive under-estimation of *C_i(c)_* when those tobacco and sunflower leaves were detached and rapidly desiccated (MS#2). There, the relative humidity in leaf airspace was estimated to be less than 80% of saturation in both species as the leaves dried up. Likewise, Cernusak et al. (2018) estimated that the relative humidity could be as low as 77% and 87% in *Pinus edulis* and *Juniperus monosperma* even when the leaves opened stomata and actively photosynthesized.

### Applications

One could do a direct *C_i_* measurement simply on one leaf side, but the greatest value is pairing with calculated values on the other side for an accurate paired measurement of water fluxes. In that case a correction of the direct measurement is needed for the vertical CO_2_ gradient especially in leaves with lower *g_ias_* (e.g., thicker leaves). Anatomical measure of *g_ias_* may be used for this correction. The dual gas-exchange system allows a robust estimation of cuticle conductance for water vapor on intact stomatous leaf surfaces in amphi-stomatous species, giving an new opportunity to study cuticular physiology in connection with stomatal physiology. The technique, owing to a high throughput (<10 min/ sample), will also allow for exploring a large number of plant species. A disadvantage is a current limitation of the approach to amphi-stomatous leaves. In hypo-stomatous leaves CO_2_ moves through the astomatous side so slowly that the closed system requires much longer times to reach equilibrium. The applicability also depends on the stomatal behavior. The estimation of *g_cw_* would be more accurate as stomatal conductance diminishes, although this is aided by simply conducting induction responses from dark-adapted leaves. In addition to *g_cw_*, location and/or unsaturation of *W_i_* deserves more investigations as both have wide implications for gas exchange and water transport in leaves.

## Acknowledgements

Thanks are due Dr. Roxana Khoshravesh for leaf anatomical measurements. We particularly appreciate thoughtful discussions with Prof. John S. Boyer (University of Missouri). This work was supported by funding to JT through the JSPS KAKENHI 18J00308 at Hiroshima University, and to DTH through the NSF EPSCoR Program under Award # IIA-1301346 and through NSF IOS 1658951 at the University of New Mexico. Any opinions, findings, and conclusions or recommendations expressed in this material are those of the authors and do not necessarily reflect the views of the National Science Foundation. JT is supported by Research Fellowships for Young Scientists from the Japanese Society for the Promotion of Science.

## Author contributions

JT planned and designed the research. JT and JRS performed experiments and analyzed data. JT and DTH wrote the manuscript.

