## Supplementary material for "Robust estimates of cuticle conductance on stomatous leaf surfaces during the light induction of photosynthesis": Fig. S1

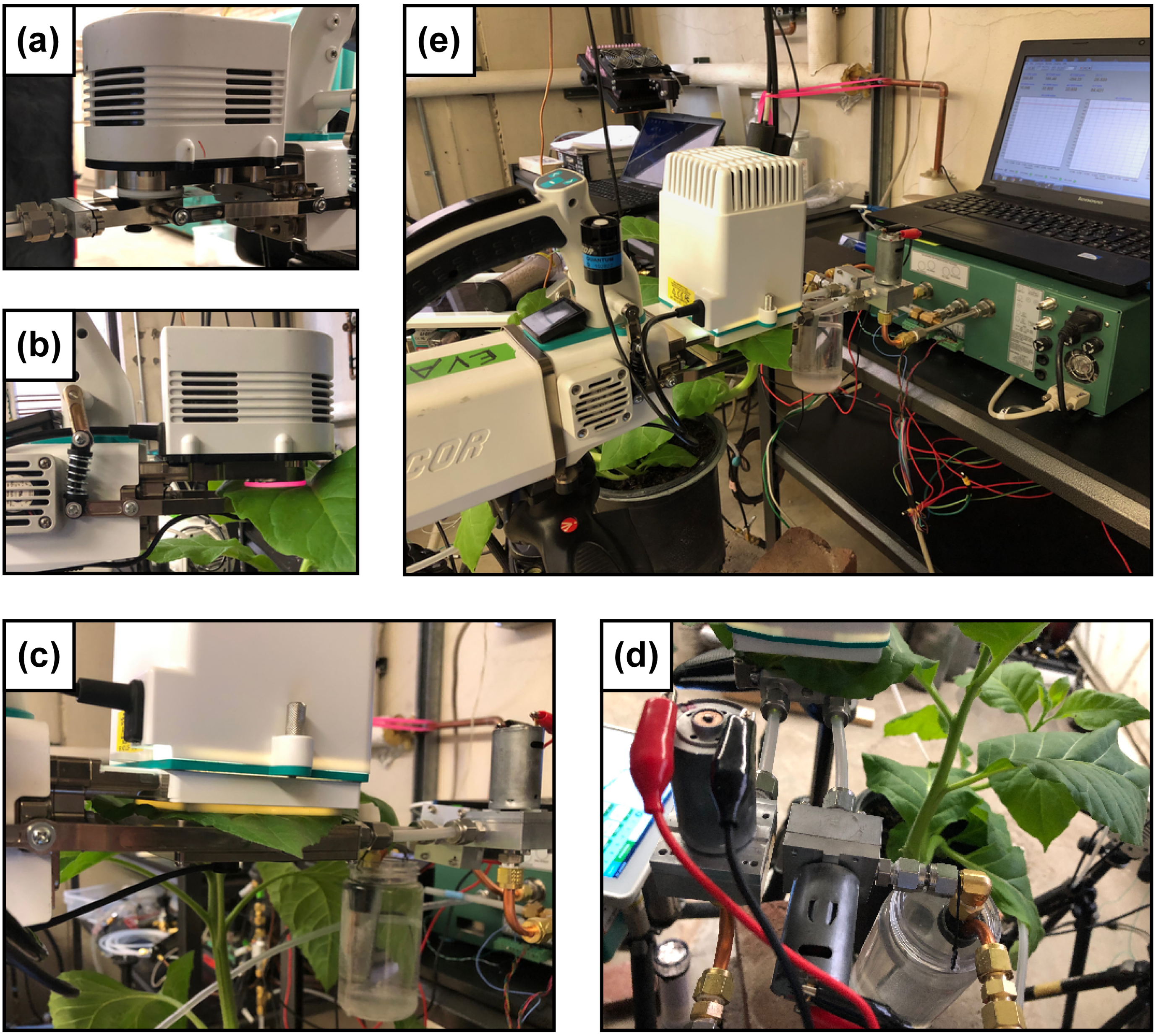


**Fig. S1.** The direct $C_i$ measurement system incorporated into a commercial open gas exchange system LI-6800 (LI-COR, Lincoln, NE, USA) equipped with a fluorometer chamber (6800-01A; LI-COR) (a and b) or a large 6 x 6 cm chamber (6800-13; LI-COR) (c, d, and e). The chamber bottom-half was rotated $180^{o}$, and attached to a solid adapter whereas the manifold to the bottom chamber was closed by a blanking plate (a). When the chamber enclosed a leaf the open system measured gas exchange only in the top chamber (i.e., adaxial side) (b and c). Two miniature impeller pumps were connected in parallel to the adapter of the rotated bottom chamber (d), and smoothly circulated the air to an infrared gas analyzer (IRGA; LI-7000; LI-COR) in a loop (e). A water jacketed glass tube (condenser) maintains the dew point slightly lower than the room temperature to prevent condensation in the IRGA (d and e).
