## Supplementary material for "Robust estimates of cuticle conductance on stomatous leaf surfaces during the light induction of photosynthesis": Fig. S2

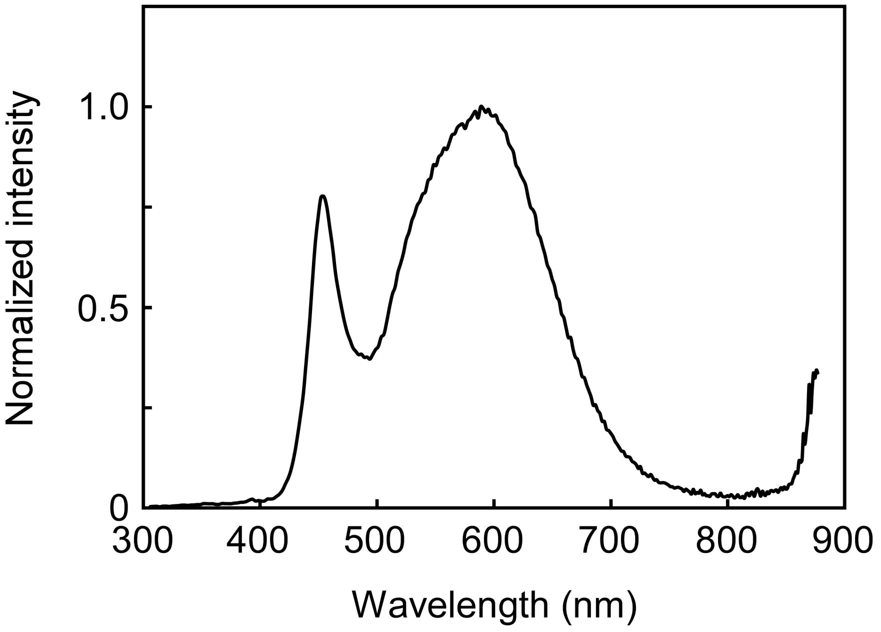


**Fig. S2.** Spectrum of the white LED of the large light source (6800-03, LI-COR) measured with a spectroradiometer (SpectraPen mini, PSI, Drasov, Czech Republic).
