## Supplementary material for "Robust estimates of cuticle conductance on stomatous leaf surfaces during the light induction of photosynthesis": Fig. S3

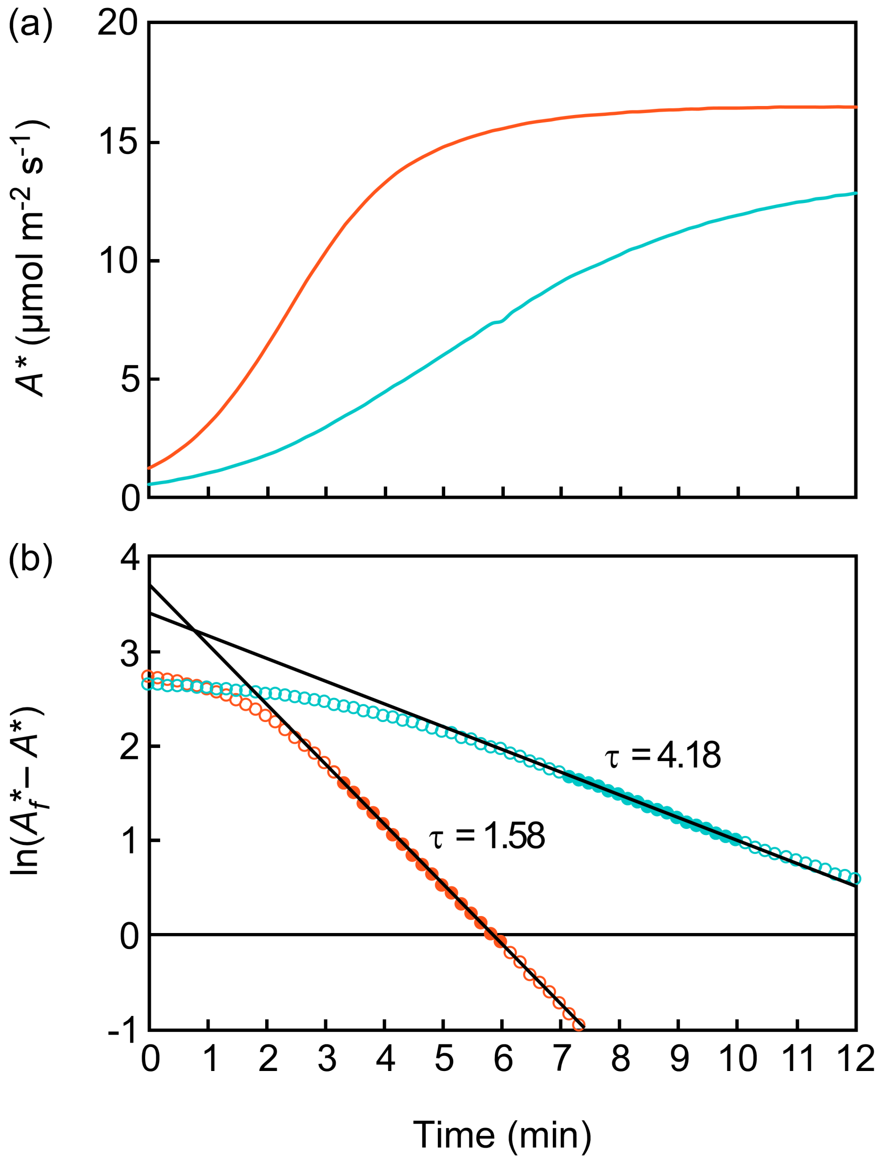


**Fig. S3.** An example of estimating the relaxation time (τ) for the rubisco phase. Time courses for the *A* normalized to $C_i$ of 250 μmol mol$^{-1}$ ($A^*$) (a), and the same data plotted as the natural logarithm of the difference between the final steady-state $A^*$ ($A_f^*$) and the $A^*$ at each time (b). Data for Fig.1 in tobacco without the pre-equilibrium is shown. In (b), the regression of the linear phase was analyzed for 3 min data (closed symbols) since $C_{i(m)}$ (orange) or $C_{i(c)}$ (blue) reached the minimum. In this example, slope values (-1/τ) were -1/1.58 ($R^2$=0.9997) and -1/4.18 ($R^2$=0.9997) with the minimum $C_{i(m)}$ of 84.4 μmol mol$^{-1}$ and the minimum $C_{i(c)}$ of 141.0 μmol mol$^{-1}$, respectively.
