## Supplementary figures and images for "Robust estimates of cuticle conductance on stomatous leaf surfaces during the light induction of photosynthesis"

### Table S1

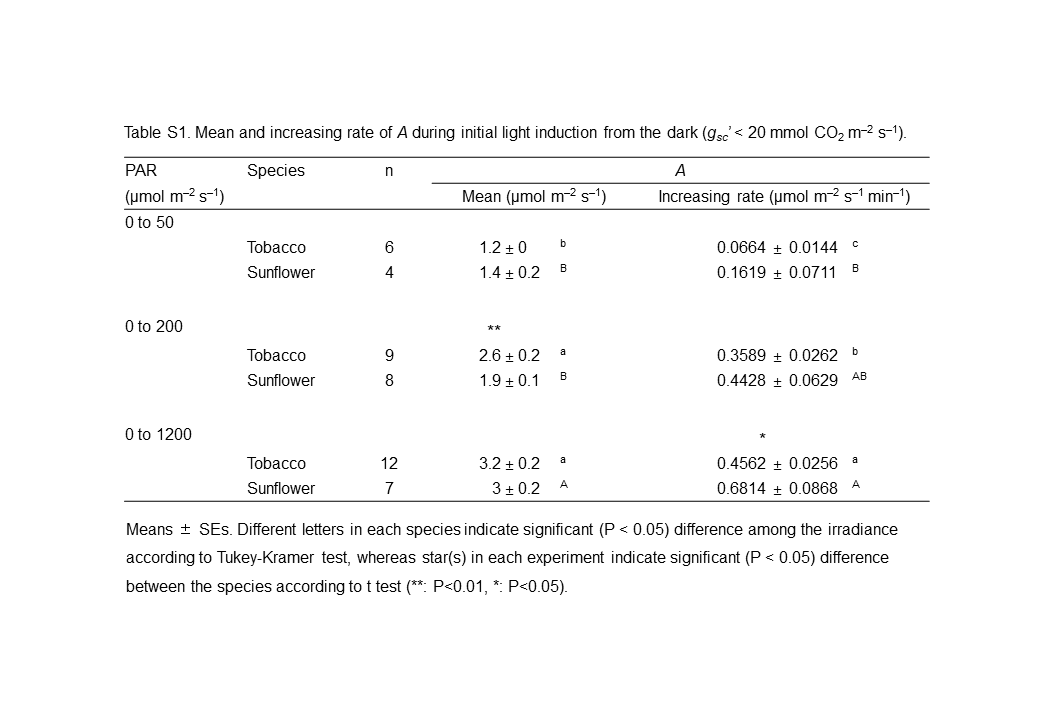
